# Japanese Encephalitis Virus NS1’ Protein regulates the expression and distribution of TIM-1 to promote viral infection

**DOI:** 10.1101/2022.12.15.520687

**Authors:** Zhenjie Liang, Shengda Xie, Junhui Pan, Xingmiao Yang, Wenlong Jiao, Ruibing Cao

## Abstract

Japanese encephalitis virus (JEV) is a mosquito-borne *Flavivirus*, which may cause severe encephalitis in humans, horses, and other animals. TIM-1 has been identified to be a receptor that promotes various viruses to enter into target cells in recent years. In the present study, we found that TIM-1 protein was significantly increased in A549 cells at the late stage of JEV infection, while the transcription levels of TIM-1 remained unaltered. Interestingly, we found that NS1’ protein plays a key role in increasing the expression of TIM-1 in cells infected with JEV. Further, we found that the NS1’ protein also efficiently regulates TIM-1 protein to distribute in the cytoplasm in JEV-infected cells, and the amount of TIM-1 protein located on the cell membrane was reduced instead. As a consequence, NS1’ protein antagonize TIM-1 mediated viral restriction for further viral infection and propagation in the late stage of infection. In molecular mechanism, the molecular weight of TIM-1 increased a bit in the present of NS1’. Expression of NEU1 down-regulated TIM-1 expression and oseltamivir treatment increased the expression of TIM-1. Therefore, our data indicated that JEV NS1’ protein facilitated the sialylation modification of TIM-1 to up-regulate the level of TIM-1 expression and regulate its distribution. Collectively, our study revealed that JEV NS1’ protein regulates the expression and distribution of TIM-1 by facilitating its sialylation to antagonize TIM-1-mediated JEV restriction for further infection. Understanding the functional interplays between TIM-1 and NS1’ proteins will offer new insights into virushost interaction.

**IMPORTANCE:** T-cell immunoglobulin and mucin domain protein 1 (TIM-1) is known to promote cellular entry of various enveloped viruses. We discovered a novel phenomenon of dynamic expression and functional regulation of TIM-1 during JEV infection. Firstly, TIM-1 protein increased in the cytoplasm at the late stage of JEV infection. Furthermore, JEV NS1’ protein up-regulated the intracellular expression of TIM-1 and antagonize TIM-1-mediated viral restriction for further viral infection. In molecular mechanism, JEV NS1’ protein facilitated the sialylation of TIM-1 to increase the level of TIM-1 expression and cytoplasm distribution. Taken together, at the late stage of infection, JEV employs a strategy in which NS1’ promotes JEV release by promoting the sialylation of TIM-1, causing it to be primarily localized in the cytoplasm. Therefore, we discovered new functions of TIM-1 and JEV NS1’ during the process of JEV infection, and provide a new insight into the interactions between JEV and cell hosts.

## INTRODUCTION

Japanese encephalitis virus (JEV) belongs to the *Flavivirus* genus of the *Flaviviridae* family, which includes more than 70 species such as West Nile virus (WNV), dengue virus (DENV), Zika virus (ZIKV) and yellow fever virus (YFV) (1, 2). JEV is the major cause of viral encephalitis in South and Southeast Asia. It leads to more than 50,000 cases in humans each year (3, 4). JEV is a mosquito-borne virus with a zoonotic transmission cycle maintained by mosquito vectors and vertebrate hosts, such as pigs and birds (5). JEV is an enveloped virus containing a single strand of positive-sense RNA approximately 11 kb in length. It encodes a single polyprotein which is proteolytically cleaved into three structural proteins (C, prM, and E) and seven nonstructural proteins (NS1, NS2A, NS2B, NS3, NS4A, NS4B, and NS5) by host and virus-encoded proteases (6). In addition, NS1’ is a novel JEV protein, which is produced through a programmed - 1 ribosomal frame-shifting mechanism (7, 8).

T cell immunoglobulin and mucin domain proteins are cell surface signaling receptors in T cells and scavenger receptors in antigen-presenting cells and kidney tubular epithelia. The human TIM protein family is composed of three members, TIM-1, TIM-3, and TIM-4 (9). TIM-1 is expressed on a broad range of epithelial cells including mucosal epithelia from the trachea, cornea, and conjunctiva tissues and is believed to be related to mosquito-borne flavivirus infection (10, 11). TIM-1 gene encodes type I cell-surface glycoproteins with a common structure including an N-terminal immunoglobulin (Ig)-like domain, a mucin domain with O-linked glycosylations and with N-linked glycosylations close to the transmembrane domain, and a cytoplasmic region with tyrosine phosphorylation and ubiquitination motif (12, 13). TIM IgV domains contain conserved residues that coordinate with metal ions such as calcium. This conserved binding pocket has been termed the metal ion-dependent ligand-binding site (MILIBS) designed for the specific recognition of phosphatidylserine (PS) (14). Phosphatidylserine receptors bind PS and mediate the uptake of apoptotic bodies. Many enveloped viruses utilize this PS receptor mechanism to adhere to and internalize into cells. This clever use of this uptake mechanism by enveloped viruses is termed apoptotic mimicry (15).

Many secreted and cell-surface expressed mammalian proteins are glycosylated, and sialic acids serve as the monosaccharide at the end of the polysaccharide chain in many of the glycosylated structures (16-18). Sialidases, named neuraminidases, remove the terminal sialic acid from these glycoconjugates (19, 20). TIM-1, as a type I transmembrane protein, has a mucin domain with O-linked glycosylation and with N-linked glycosylation, also occurring multiple other modifications including alpha1-(3,4)-fucosylation, tyrosine sulfation, and so on (21).

JEV NS1 is a multifunctional non-structural protein involved in virus replication and regulation of host innate immunity. JEV NS1’ protein is the product of a −1 ribosomal frameshift event that occurs at a conserved slippery hexanucleotide motif located near the beginning of the NS2A gene and is stimulated by a downstream RNA pseudoknot structure which contains NS1 protein and subsequent 52 amino acids (22). It is reported that NS1’ is related to the neuro-invasiveness of JEV (8). WNV NS1’ can completely replace the key role of NS1 in the virus life cycle in cells (23). JEV NS1’ facilitates virus proliferation in avian cells and chicken embryo (24). Recently, it has been proved that NS1’ protein interrupts the CDC25C phosphatase-mediated dephosphorylation of CDK1, which prolongs the phosphorylation status of CDK1 and leads to the inhibition of MAVS-mediated IFN-β induction. However, more functions of NS1’ still need to be explored. (25, 26).

In this study, we found that NS1’ protein increased the TIM-1 expression on protein level in a late stage of JEV infection. We also found that NS1’ regulated the distribution of TIM-1 to counteract TIM-1-mediated inhibition of JEV release for further infection and transmission. In addition, our results indicated that NS1’ protein promoted TIM-1 cytoplasmic distribution by facilitating its sialylation, making it more stable and non-degradable. Taken together, this study enriched the function of JEV NS1’ and provided some new ideas for more functions of TIM-1.

## RESULTS

### JEV Infection increases TIM-1 expression in A549 and Vero cells

In previous experiments, we have found that the expression of TIM-1 protein in cells can promote JEV infection. By chance, we discovered that TIM-1 expression in A549 cells had a dramatic increase following JEV infection. To ensure this, the A549 cells were infected with the JEV strain NJ08 at a multiplicity of infection (MOI) of 1 and collected samples at different times after infection. Western blot analysis and immunofluorescence staining results both indicated that the level of TIM-1 protein was significantly up-regulated after JEV strain NJ08 infection (Fig.1A, 1B). The increased TIM-1 expression was related to the inoculated JEV dose (Fig.1C). However, mRNA levels of TIM-1 were unaltered following JEV infection (Fig.1D), suggesting that posttranscriptional modification of TIM-1 may be occurred. What’s more, this phenomenon was elevated in Vero cells, the results also indicated that the expression of TIM-1 was significantly increased. (Fig.1E). Taken together, these findings demonstrated that TIM-1 expression in cells was up-regulated both in time-dependent and JEV dose-dependent manners at protein levels in cells with JEV infection.

**Figure 1.**
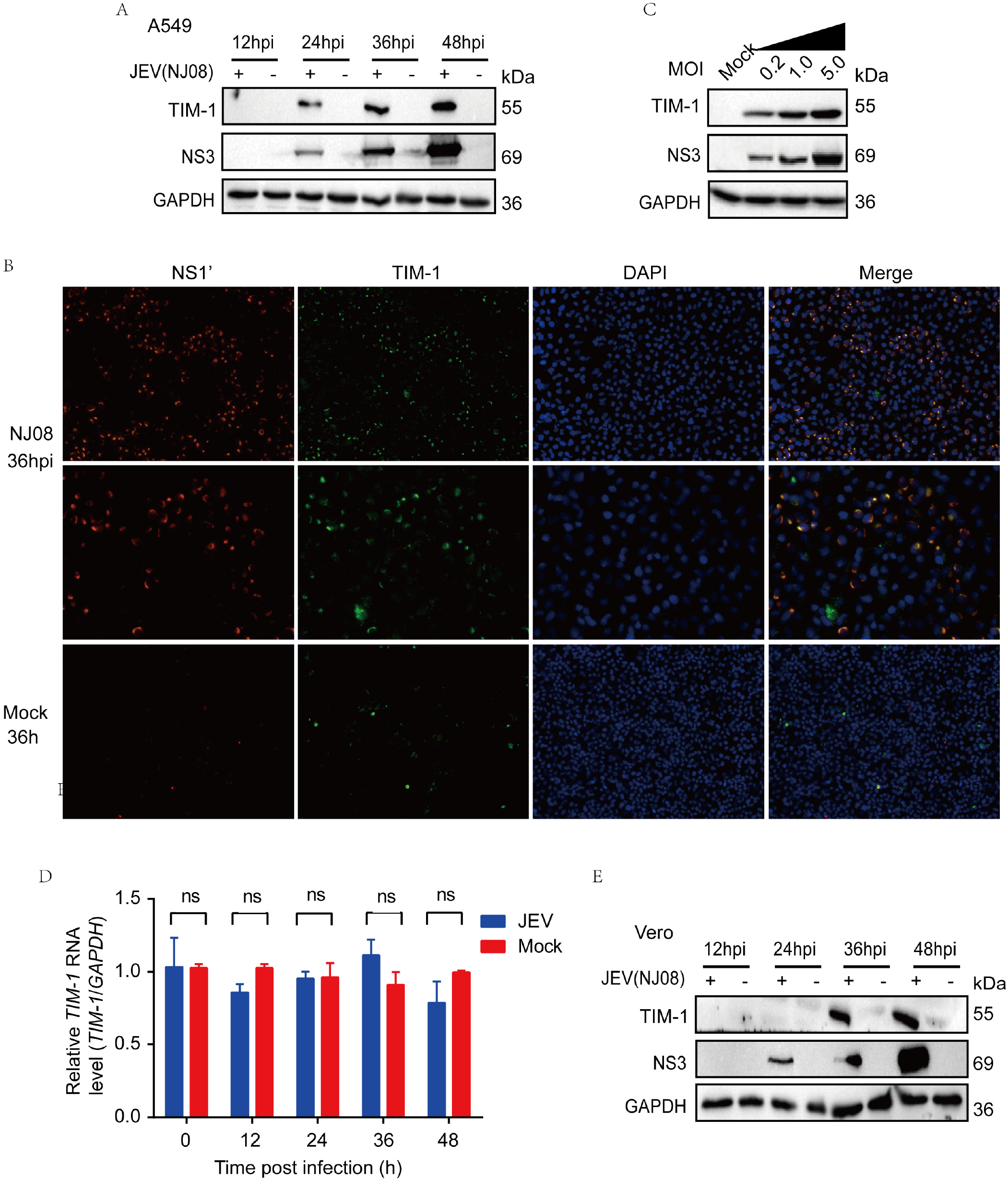
JEV infection increases TIM-1 protein expression in cells. A, A549 cells were challenged with JEV strain NJ08 at MOI= 1 and collected samples at various time (12hpi、24hpi、 36hpi、48hpi) for western blot analysis. B, A549 cells were infected with JEV strain NJ08 and fixed at 36hpi. Immunofluorescent (IF) staining with TIM-1 (green) and JEV-E (red) and the nuclei with DAPI (4’,6-diamidino-2-phenylindole). C, A549 cells were challenged with JEV strain NJ08 for 36hpi at different MOI for western blot analysis. D, A549 cells were challenged with JEV NJ08 strain, TIM-1, E, and GAPDH mRNA levels were determined by real-time quantitative reverse transcription polymerase chain reaction (qRT-PCR). Data are presented as means ± SD, experiments are run with duplicate samples. ns, not significant. E, Vero cells were challenged with JEV NJ08 strain at different points, and JEV NS3 and GAPDH were evaluated.

### NS1’ protein plays key role in increasing the expression of TIM-1 in cells infected with JEV

To investigate the mechanisms through which JEV infection up-regulated TIM-1 expression, we initially determined whether JEV-encoded proteins were related to TIM-1 expression. Three plasmids encoding structural proteins of JEV (C, M, and prME) and six plasmids encoding nonstructural proteins of JEV (NS1’, NS1, NS2B, NS3, NS4B, and NS5) were transfected into A549 cells, separately. Interestingly, NS1’ protein, rather than other JEV proteins increased the expression of TIM-1 protein (Fig.2B). And TIM-1 expression was significantly up-regulated by NS1’ in a dose-dependent manner (Fig.2C). Similarly, immunofluorescence staining confirmed that JEV NS1’ protein remarkably elevated the expression of TIM-1 in cells, while there was no significant change on TIM-1 mRNA level at indicated time points (Fig. 2D-2E). At the same time, we also tested whether the recombinant NS1’ protein could increase the expression of TIM-1 protein. As shown in Fig. 2F and 2G, the expression of TIM-1 was also increased in cells after recombinant NS1’ protein treatment. Considering that JEV attenuated vaccine strain SA14-14-2 does not express NS1’ protein, while JEV virulent isolates all express NS1’. We inoculated A549 cells with gene type I JEV strain HN07, gene type III JEV strain NJ08, and JEV attenuated vaccine strain SA14-14-2, respectively. And then analyzed the expression of TIM-1 protein in cells by Western blot. As expected, TIM-1 expression significantly increased in A549 cells infected with JEV strains HN07 and NJ08, while there was no obvious increase expression of TIM-1 protein in JEV strain SA14-14-2 inoculated cells (Fig.2H). Collectively, these findings indicated that JEV NS1’ protein mediated the increased expression of TIM-1 protein in cells infected with JEV.

**Figure 2.**
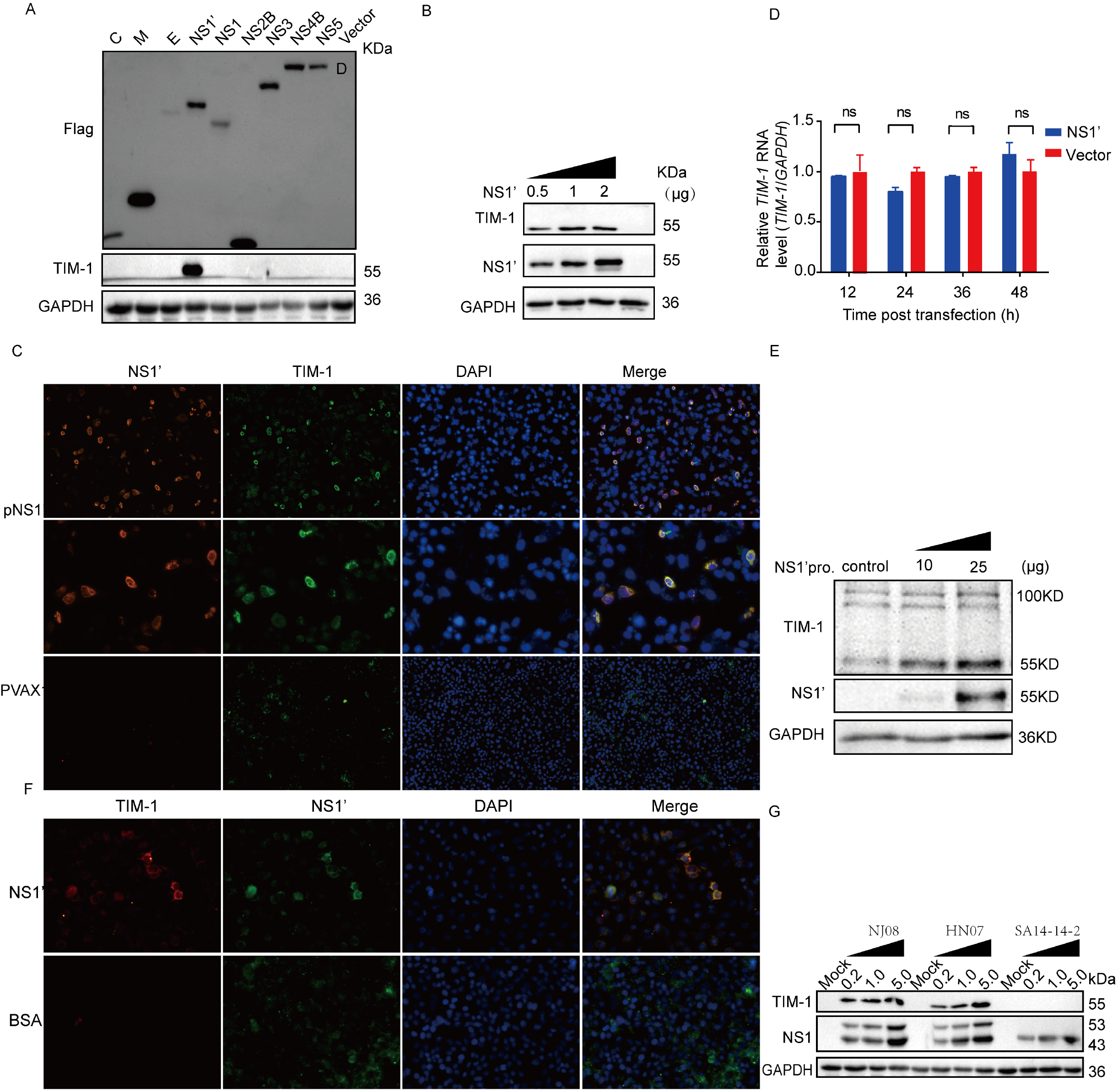
TIM-1 protein expression was up-regulated by JEV NS1’. A, A549 cells were challenged with JEV strain NJ08, HN07, and SA14-14-2 at MOI 0f 0.2, 1, and 5, the cells were lysed at 36hpi for western blot analysis. B, A549 cells were transfected with C, -M, -E, -NS1’, - NS1, -NS2B, -NS3, -NS4B, or -NS5 plasmids for 36h. TIM-1, C, M, E, NS1’, NS1, NS2B, NS3, NS4B, NS5, and GAPDH proteins were determined by western blotting with the indicated antibodies. C, A549 cells were transfected with p3XFLAG-CMV-7.1-NS1’ using a concentration gradient for 36h. TIM-1, NS1’, and GAPDH proteins were determined by western blotting. D, A549 cells were transfected with pVAX1-NS1’ for 36h and then fixed with 4% paraformaldehyde. Immunofluorescent (IF) staining with TIM-1 (green) and JEV-NS1’(red) and the nuclei with DAPI (4’,6-diamidino-2-phenylindole). E, A549 cells were transfected with pVAX1-NS1’, and TIM-1 mRNA levels were measured by qPCR. Data are expressed as means ± SEMs of three independent experiments. ns, not significant. F, Adding the indicated concentration of purified NS1’ protein into A549 cells, the lysates were collected after incubating for 24h and analyzed using western blot. G, Adding the indicated concentration of purified NS1’ protein into A549 cells, Immunofluorescent (IF) staining with TIM-1 (green), and JEV-NS1’(red) and the nuclei with DAPI (4′,6-diamidino-2-phenylindole). H, 293T cells were co-transfected with TIM-1 and C, -M, -E, -NS1’, -NS1, -NS2B, -NS3, -NS4B or -NS5 plasmids, respectively. After 48h post transfection, total TIM-1, NS1’, and GAPDH proteins expressed in cells were detected by western blotting with the indicated antibodies. I, 293T cells were co-transfected with TIM-1 and NS1’ for 36h and then fixed using 4% paraformaldehyde. Immunofluorescent (IF) staining with TIM-1 (green) and JEV-NS1’ (red) and the nuclei with DAPI (4′,6-diamidino-2-phenylindole).

### TIM-1 protein in JEV-infected cells mainly distributed in the cytoplasm

Considering that TIM-1 can promote virus-cell adsorption, we explored whether the increased TIM-1 was primarily located on the cell surface. JEV strain NJ08 was inoculated to A549 cells, then the immunofluorescence assay was carried out with one group permeabilized and another as the control. The results showed that increased TIM-1 protein mainly distributed in the cytoplasm after JEV infection in cells permeabilized with 0.1% triton-100, and TIM-1 is co-localized with NS1’ (Fig.3B). Whereas TIM-1 protein specific fluorescence did not change significantly in JEV infected cells and control cells under the non-permeabilization condition (Fig.3A). This result is inconsistent with our expectation. After JEV infection, TIM-1 expression on the cell surface did not increase, or even decreased. The up-regulated TIM-1 was mainly located in the cytoplasm. Similarly, A549 cells were transfected with pNS1’, TIM-1 specific fluorescence increased significantly only occurred in the permeabilization group (Fig.3D), whereas TIM-1 specific fluorescence in the control group showed no apparent difference (Fig.3C). Consequently, we preliminary concluded that increased expression of TIM-1 protein in JEV infected cells is mainly distributed in the cytoplasm.

**Figure 3.**
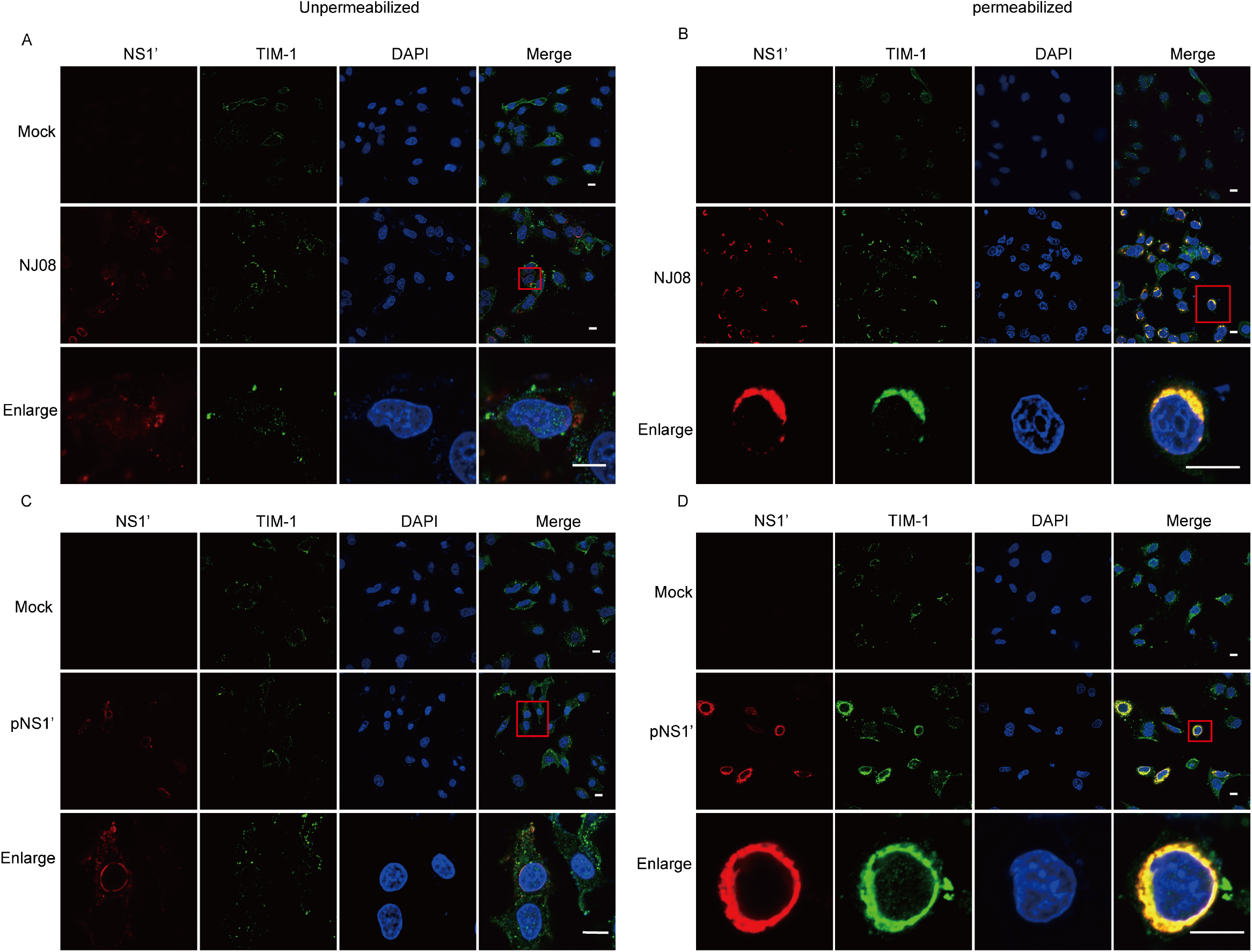
Increase expressed TIM-1 protein is mainly distributed in the cytoplasm. A-B, The A549 cells were challenged with JEV strain NJ08 at MOI=0.1 for 36 hours, and then the cells were fixed with 4% paraformaldehyde. According to the Confocal fluorescence Microscopy assay, Immunofluorescent (IF) staining with TIM-1 (green), JEV-NS1’ (red), and the nuclei with DAPI (4’, 6-diamidino-2-phenylindole), the right group was permeabilized, the left group was unpermeabilized, scale bars, 10 μm. C-D, The A549 cells were transfected with pNS1’ for 36 h and then fixed. The experimental method was the same as above A and B, scale bars, 10 μm.

### NS1’ protein antagonizes TIM-1–Mediated Restriction of JEV

In the early stage of JEV infection, it is clear that TIM-1 acts as a co-receptor that promotes the viral infection. However, the biological significance of the increased TIM-1 in the cytoplasm caused by NS1’ protein at the late stage remains unknown.

It has recently been demonstrated that TIM-1 can inhibit HIV release and HIV Nef regulates the cellular distribution of TIM-1 to antagonize TIM-1-mediated HIV restrcition to enhance viral infection (27). We hypothesize that the role of NS1’ protein is similar to HIV Nef protein at the late stage of infection. To test this hypothesis, A549 cells were infected with a pair of JEV strains differently expressing NS1’ protein including vaccine strain SA14-14-2 (without NS1’ protein expression) and its mutant strain A66G (expressing NS1’ protein) at the MOI of 0.1 and collected the cell and supernatant samples at indicated times for western blot and qPCR, separately. The results showed that the infection of JEV strain A66G in A549 cells was higher than that of JEV strain SA14-14-2 throughout the infection phase. However, the virus in A549 cells infected with JEV strain SA14-14-2 were higher than that of JEV strain A66G at the late stage of infection, whereas the amount of virus in the supernatant were on the contrary (Fig.4A-4B). This result leads us to believe that the role of the JEV NS1’ protein is to antagonize TIM-1-mediated JEV restriction as the HIV Nef. Therefore, we may conclude that the NS1’ protein is capable of efficiently regulating the expression and distribution of TIM-1 and antagonizing the adhesion between TIM-1 and progeny viruses for virus release and further transmission in the late stage of infection.

In addition, 293T cells transfected with plasmids of TIM-1 and/or NS1’ with different doses were infected with the JEV strain SA14-14-2 at an MOI of 1 for indicated hours. The cells transfected with TIM-1 did show a dose-dependent enhancement of JEV infection, while a graded decrease in infection was observed in cells co-transfected with different concentrations of pNS1’ and pTIM-1 (Fig.4C-4D). This phenomenon aroused our concern. NS1’, as a viral protein, played an inhibitory role in JEV infection. We considered that there was an essential difference between the production time of NS1’ in this experimental model and that of natural infection. When the A549 cells were co-transfected with TIM-1 and NS1’, TIM-1 protein mainly distributed in the cytoplasm in the early stage and cannot play the role of a receptor, which reversely proved that JEV NS1’ mainly promotes TIM-1 cytoplasmic location to limit its adhesion to virus in the late stage of infection. Therefore, these results elucidated that TIM-1 plays dual roles of promoting viral entry at an early stage and inhibiting the viral release at the late stage, and JEV evolved different strategies to cope with it correspondingly.

### JEV NS1’ protein regulates TIM-1 expression and distribution by promoting its sialylation

Having identified that JEV NS1’ could up-regulate the expression of TIM-1 protein, we continued to explore whether NS1’ inhibited the proteasome degradation of TIM-1. After treated cells with the proteasome inhibitor MG132, TIM-1 expression was up-regulated. However, an interesting phenomenon exists: in western-blot results, the TIM-1 bands from TIM-1 and NS1’ co-transfected group are slightly larger than those in cells transfected TIM-1 alone (Fig. 5A), in line with the results of Fig. 4c-4d. The molecular weight of the CD45 monomer becomes slightly larger after salivation modification, and it cannot form dimer and cellular location changed, which affected its function (28). This phenomenon is similar to our results. Therefore, we wondered if NS1’ could regulate the sialic acid modification of TIM-1. To verify this hypothesis, we co-expressed NEU1 and NS1’ in A549 cells, the western-blot results indicated that the amount of 55 KD band of TIM-1 was decreased while the 100KD band was increased compared with the group transfected NS1’ plasmid alone (Fig. 5B). In addition, we co-transfected pNEU1, pNS1’ and pTIM-1 into 293T cells, the results showed that NEU1 could counteract NS1’-mediated the up-regulation expression of TIM-1 at 55KD form. (Fig. 5C). Oseltamivir phosphate, an inhibitor of NEU1, was added to A549 and 293T cells transfected with TIM-1. The expression of the 55KD form of TIM-1 was increased, which is similar to the role of NS1’ protein (Fig. 5D, 5E). The optimal Oseltamivir dose has no obvious effects on cell viability (Fig. 5F, 5G). Collectively, JEV NS1’ is thought to regulate the expression and distribution of TIM-1 via facilitating its sialylation to enhance JEV infection at the late stage.

**Figure 4.**
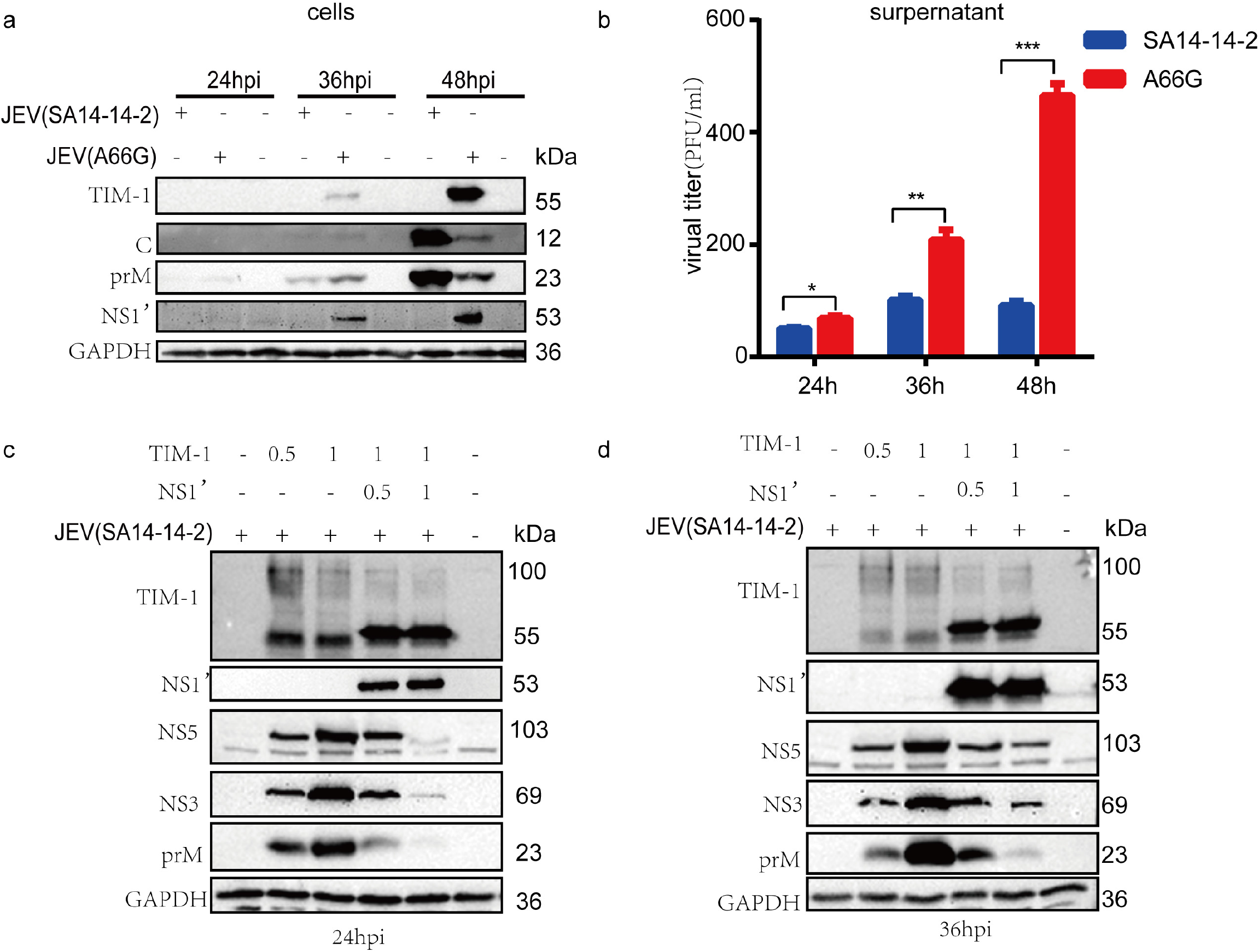
NS1’ protein antagonizes TIM-1–Mediated Restriction of Virus. A, A549 cells were inoculated with JEV strain SA14-14-2 and JEV strain SA14-14-2 NS2A A66G, at 24, 36, and 48 post transfection, cells and supernatant were collected respectively. A representative Western blot analysis is demonstrated. The expression of TIM-1, JEV C, prM, and GAPDH was assessed. B, A549 cells were challenged with JEV strain SA14-14-2 and strain A66G, and the virus in the supernatant was determined by real-time quantitative reverse transcription polymerase chain reaction (qRT-PCR). Data are presented as means ± SD of n = 3 experiments run with duplicate samples. C-D, HEK293T cells were transfected with TIM-1 and NS1’ plasmids. 24 hours post-transfection, cells were inoculated with JEV strain SA14-14-2 for 24/36 hpi. Western blotting was performed to measure viral proteins including structural protein prM and non-structural protein NS3 NS5. TIM-1 and NS1’expression in cell lysates were determined by using an anti–TIM-1 and anti-NS1’ antibodies, respectively.

**Figure 5.**
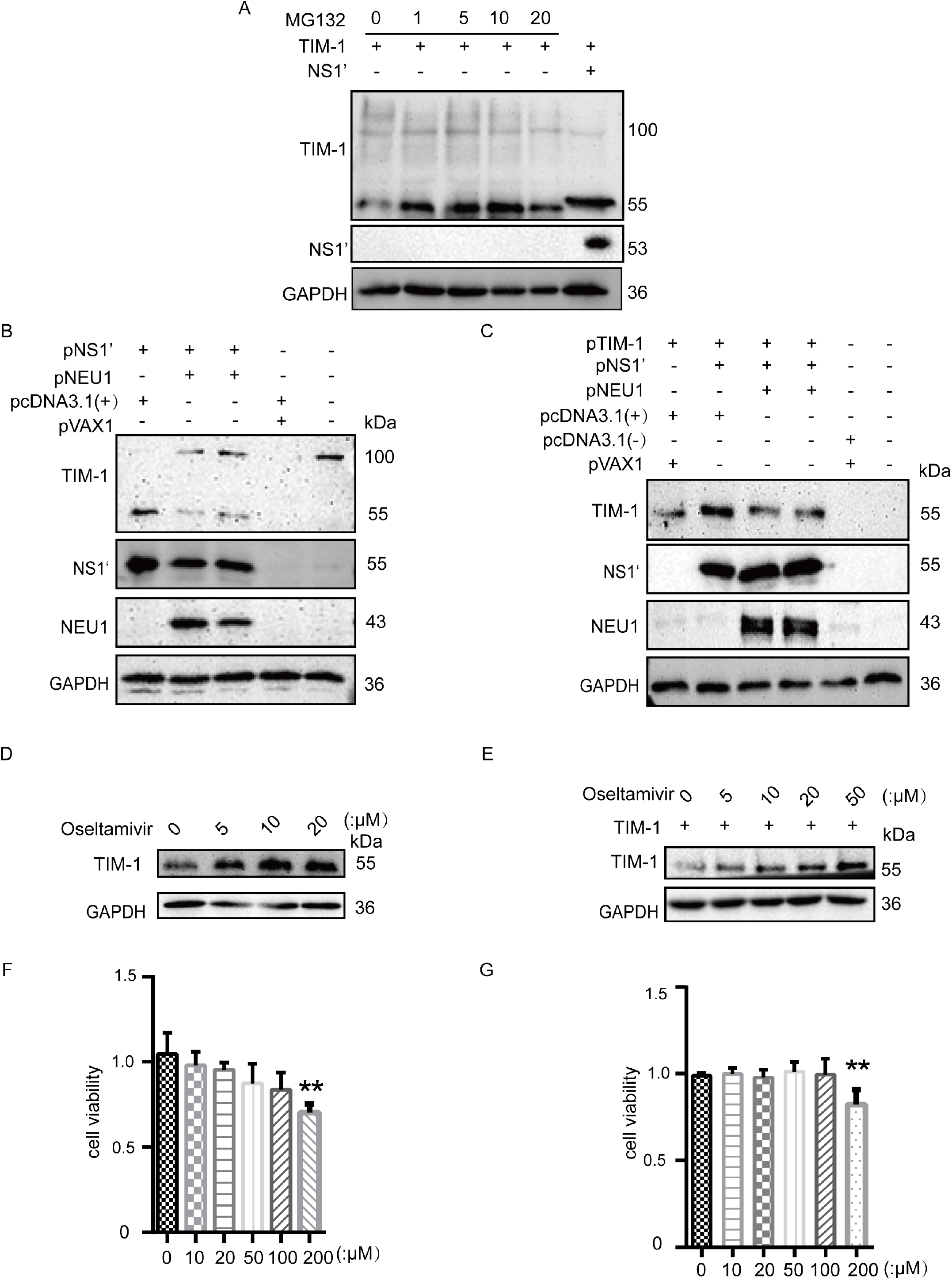
JEV NS1’ increases the expression of TIM-1 by promoting its sialylation. A, HEK293T cells were transfected with plasmid pcDNA3.1(+)-TIM-1 or pVAX1-NS1’. 24h later, cells were treated with different amount of proteasome inhibitor MG132 (1μM,5μM,10μM,20μM), and another 24h later cells were lysed and subjected to western blot with indicated antibodies. B, A549 cells were co-transfected with pcDNA3.1(+)-NEU1 and pVAX1-NS1’, total TIM-1, NS1’, NEU1, and GAPDH proteins expressed in samples were detected by western blotting with the indicated antibodies. C, 293T cells were co-transfected with pcDNA3.1(-)-TIM-1、pcDNA3.1(+)-NEU1 and pVAX1-NS1’, TIM-1, NS1’, NEU1 and GAPDH proteins expressed in cells were detected by western blotting with the indicated antibodies. D, A549 cells were treated with Oseltamivir phosphate for 24h, and TIM-1 proteins were detected by western blotting with the indicated antibodies. E, 293T cells were transfected with TIM-1 plasmid for 24h, then were treated with indicated Oseltamivir phosphate for another 24h, TIM-1 proteins were detected by western blotting. F-G, Adding the CCK-8 (10 μ L) to the A549 or 293T cells for 1.5h, the results were measured by microplate reader at 450nm.

## DISCUSSION

We have previously proved that TIM-1 can promote JEV entry cells and infection (29). In the present study, we found the expression of TIM-1 protein had a significant increase in A549 and Vero at the late stage of JEV infection. The qPCR results showed that there was no significant difference in the transcription level of TIM-1 at different time points. Then we confirmed that JEV NS1’ mediated the increase of TIM-1 at the protein level. And immunofluorescence results revealed that the increased TIM-1 regulated by JEV NS1’ is mainly distributed in the cytoplasm. These results suggested that NS1’ protein changed the expression and distribution to abolish the TIM-1-mediated constraint of JEV for further transmission. In addition, overexpression of NS1’ in A549 cells, oseltamivir phosphate inhibition assays, and fluorescence staining showed that JEV NS1’ protein up-regulated the overall level of TIM-1 expression and promoted TIM-1 intracellular location via facilitating its sialylation. Taken together, we concluded that JEV NS1’ regulated the expression and distribution to promote JEV infection.

TIM-1, an immune co-stimulatory molecule, is expressed at a higher level in tumor tissues than in normal tissues in a variety of cancers, including cervical cancer, gastric cancer, lung cancer, glioma, and others (30-32). Higher TIM-1 expression in cancer tissues may be used as an independent prognostic predictor for patients. Increased expression of TIM-1 is also linked to T cell immune tolerance (33, 34). Therefore, the increased expression of TIM-1 is the result of organismal or viral regulation in the specific situation.

TIM-1 acted as virus attachment or entry cofactors, promoting infection of various enveloped viruses at an early stage (29, 35-39). It is thought to occur through the PS-binding activity of the TIM-family proteins (40). However, the function of TIM-1 is not limited to this. Li et al. have proved that TIM-1 can inhibit the release of HIV from cells related to the PS binding (41). They also found that HIV-1 Nef not only up-regulates the overall level of TIM-1 expression but also sequesters TIM-1 in intracellular compartments to abolish this restriction, which is similar with our findings (27). The AXL protein, belonging to the PS receptor, also inhibits the release of JEV. JEV NS2B-3 protein complex promotes viral particle release by down-regulating the expression of AXL (42). In our study, A549 cells were infected with a pair of JEV strains differently expressing NS1’ protein at low MOI to reduce the influence of NS1’ protein in the supernatant. Our results indicated that NS1’ antagonized the TIM-1-mediated JEV restriction via promoting TIM-1 distributing in the cytoplasm.

Combining the above-mentioned results, we speculated that partial TIM-1 is distributed on the cell surface as the form of a dimer (100KD) in its natural state in A549 cells. The monomeric form of TIM-1 protein was increased in A549 cells transfected with pVAX1-NS1’, whereas the dimeric form was upregulated in cells co-transfected with pVAX1-NS1’ and pcDNA3.1-NEU1. Therefore, we considered that NS1’ protein regulates the level of TIM-1 expression as well as its distribution. The CD45 can’t form a dimer form and localized correctly after sialylation modification, which affects its function [34]. This phenomenon is similar to our experimental results (Fig.5A). Overexpressed pNEU1 in A549 cells resulted in a more dimeric form and a less monomeric form of TIM-1. What more, after oseltamivir phosphate, a sialidase inhibitor, was added to the cells and the expression of TIM-1 was increased in the form of 55KD, just as the effect of NS1’ protein. We then had an attempt to enrich the increased expressed TIM-1 using IP assays and then treated with NEU1 protein to detect the molecular weight of the TIM-1 band, but we did not have the desired result. We speculate that it is due to the sialylation of TIM-1 may be altered during the enrichment process by some reagent treatment. In all, we concluded that NS1’ changed TIM-1 expression and distribution by facilitating its sialylation to favor virus infection.

According to our findings, after sialylation, the monomer form of TIM-1 was mostly located in the cytoplasm, with less dimer form TIM-1 on the cell surface. One perspective could explain this phenomenon: sialic acid on the TIM-1 can mask the mucin-typed sugar chain, preventing glycosidase hydrolysis and protein degradation(43-46). And the negative charge repulsion between the sialic proteins makes the molecules more stretchable, potentially preventing the proteins from forming dimeric form(47, 48). The TIM-1 protein degradation pathway is primarily via the ubiquitin-protease pathway(49). When the proteasome inhibitor MG132 was added into the control group, the TIM-1 expression increased, demonstrating that the ubiquitination degradation pathway of TIM-1 protein was inhibited. Further, the vesicles and related transported endosome may be altered and TIM-1 protein was retained in the cytoplasm. The specific reasons require further investigation.

Taken together, TIM-1 protein plays different roles at different times in the JEV life cycle, and the viruses use different strategies to deal with it accordingly. JEV NS1’ protein regulates the expression and distribution of TIM-1 to promote its infection at a late stage of JEV infection. Understanding the functional interplays between TIM or PS receptors and NS1’ proteins will offer new insights into virus-host interaction.

## MATERIALS AND METHODS

### Cells and viruses

Human embryonic kidney cells (HEK-293T), baby hamster kidney cells (BHK-21), African green monkey kidney cells (Vero), and human cervical cancer cells (HeLa) were maintained in Dulbecco’s modified Eagle’s (DMEM, 11965092, GIBCO, Carlsbad, CA, USA) medium, supplemented with 10% fetal bovine serum (FBS, 10100, GIBCO) and100 μg/ml streptomycin, and 100 IU/ml penicillin (Gibco, Invitrogen) at 37 °C with 5% CO_2_. Lung adenocarcinoma cell line (A549) and Aedes albopictus cell line (C6/36) were grown in Roswell Park Memorial Institute 1640 (RPMI 1640, 31870082, GIBCO) medium with 10% FBS and 100 μg/ml streptomycin, and 100 IU/ml penicillin (Gibco, Invitrogen). The JEV strains NJ2008 (GenBank: GQ918133.2), HN07 (GenBank: FJ495189.1), and SA14-14-2 (GenBank: MK585066.1) were preserved in our laboratory. The titers of the virus were titrated on BHK-21 cells by plaque assays.

### Plasmids, antibodies, and chemical inhibitors

The human TIM-1 and JEV NS1’ genes were cloned into pcDNA3.1 (-) and pVAX1, respectively. The capsid, prM, envelope, NS1, NS1’, NS2B, NS3, NS4B, NS5 genes of JEV strain NJ2008 were inserted into p3×FLAG-CMV-7.1. Sialidase 1 (NEU1) was cloned into pcDNA3.1 (-). Specific monoclonal antibodies for JEV NS1’ protein and NS1 protein were produced in our lab. Anti-NS3 monoclonal antibody (GTX125868), anti-E (GTX125867) polyclonal antibody, anti-Capsid (GTX131368) polyclonal antibody, and anti-prM (GTX131833) polyclonal antibody were purchased from GeneTex (Irvine, CA, USA). Anti-Human TIM-1 monoclonal antibody (MAB1750) and polyclonal antibody (AF1750) were purchased from R&D systems (Minnesota, USA). Anti-GAPDH polyclonal antibody (sc-25778) and NEU1 monoclonal antibody (sc-166824) were purchased from Santa Cruz (Dallas, USA). Anti-FLAG-tag mouse monoclonal antibody (66008-3-Ig) was purchased from protein tech (Wuhan, China). MG132 and Oseltamivir phosphate (abs47047649) were purchased from Absin (Shanghai, China). Donkey antigoat IgG-Alexa Fluor488 (abs20026), donkey anti-Rabbit IgG Alexa Flour 594 (abs20021), and Donkey anti-mouse Alexa Flour647 (ab150107) were purchased from Abcam (Cambridge, UK).

### Western blot

Confluent cell monolayers of A549 cells and 293T cells were seeded in plates and then transfected plasmids or infected with JEV. After indicated hours, cells were rinsed with cold PBS three times, then monolayers were lysed using RIPA buffer (89900, ThermoFisher, Waltham, USA) for 15 min at 4 °C. Cell lysates were separated by 10%−12.5% gradient SDS-PAGE using a running buffer at a constant voltage of 120V for 1.5 hours. For subsequent analyses, proteins were transferred onto PDVF membranes using a Trans-Blot® Turbo Transfer System (BioRad). Then, membranes were blocked in PBST with 5% skimmed milk for 1.5 hours at room temperature. Primary antibodies were incubated overnight at 4°C. Secondary antibodies were used to detect the expression of proteins.

### Quantitative Real-time Polymerase Chain Reaction (qPCR)

The total RNA in cells was extracted by the Total RNA extract KIT (OMEGA, Bio-tech, Norcross, GA, USA). Then were reverse-transcribed into cDNA using PrimeScript RT Master Mix Kit (Takara). qPCR was performed by TB Green Premix Ex Taq (Takara, RR420Q,), with gene-specific primer pairs (Table S1). The quantification of the gene expression was calculated according to the 2-ΔΔCt method.

### Virus infection

A549 or others were seeded in plates (12-well), incubated with an appropriate multiplicity of infection of JEV strain NJ08, HN07, or SA14-14-2 at 37°C for 1.5 h. Then they were washed by PBS three times and cultured with 2% FBS fresh medium at 37°C for a specified period of time. The supernatants or cell lysates were collected and applied to western blot analysis.

### Inhibitor treatment

Oseltamivir phosphate was dissolved in ddH_2_O at a suitable concentration and MG132 was melted in DMSO. 293T cells were co-transfected with TIM-1 and NS1’ plasmids, and A549 cells transfected with NS1’. And the cells were treated with prepared oseltamivir phosphate or MG132. 24 hours after transfection, the cell lysates were then subjected to western blot analysis.

### Immunofluorescence assays (IFA)

A549 cells were seeded in a 3.5 cm glass-bottom dish with a density of 2×10^5^ cells/well and incubated at 37°C with 5% CO_2_ overnight. Cells were transfected with plasmids or infected with JEV. After 24-48 hours, the cells were washed with PBS and fixed with 4% paraformaldehyde at RT for 15 min. And then cells were permeabilized with 0.1% Triton X-100 at RT for 10 min. The penetrated cells were soaked with 2% BSA to block on a nutator at RT for 0.5-1h. The cells were then incubated with a primary antibody at 4 °C overnight. The next day after washing, the cells were incubated with mixed second antibodies at 37 °C for 45 mins. At last, Nuclei were stained using 4,6-diamidino-2-phenylindole (DAPI) for 5 mins. Images were captured and analyzed by fluorescence microscopy or confocal microscopy.

### Plaque Assay

BHK-21 cells were seeded in 6-well plates (3×10^5^ cells/well) and then incubated at 37°C with 5% CO_2_. Cells were inoculated with JEV in 10-fold serial dilution using serum-free DMEM and incubated at 37°C with 5% CO_2_ for 1.5 h. After removal of virus suspension, the cells were overlaid with a 1:1 mixture of 2% low-melting agarose and 2-fold DMEM including 4% FBS and 1% P/S. When the plaque appeared in the plate, the cells were stained with crystal violet for 4 h. Finally, the titer of the virus was determined by counting the number of the virus plaques.

### Cell viability assay

Cell viability was determined using the CCK8 kit. A549 and 293T cells were treated with different compounds and incubated at 37°, 5% CO_2_, and then inoculated with 10 μl CCK8. 1.5 h later, the mixture was determined by measuring the absorbance value at 450 nm.

### Statistical analysis

Graphical representation and statistical analyses were performed using Prism6 software (GraphPad). A T-test was used to compare the data from the pairs of treated and untreated groups. Results are shown as means ± SD from three independent experiments. Statistical significance was indicated by asterisks (*, P < 0.05; **, P < 0.01) in the figures.

## ACKNOWLEDGEMENTS

This work was carried out with the support of grants from the National Natural Science Foundation of China (Grant No. 32273040) and the National Key Research and Development Plan of China (Grant No. 2022YFD1800100).

## LEGENDS

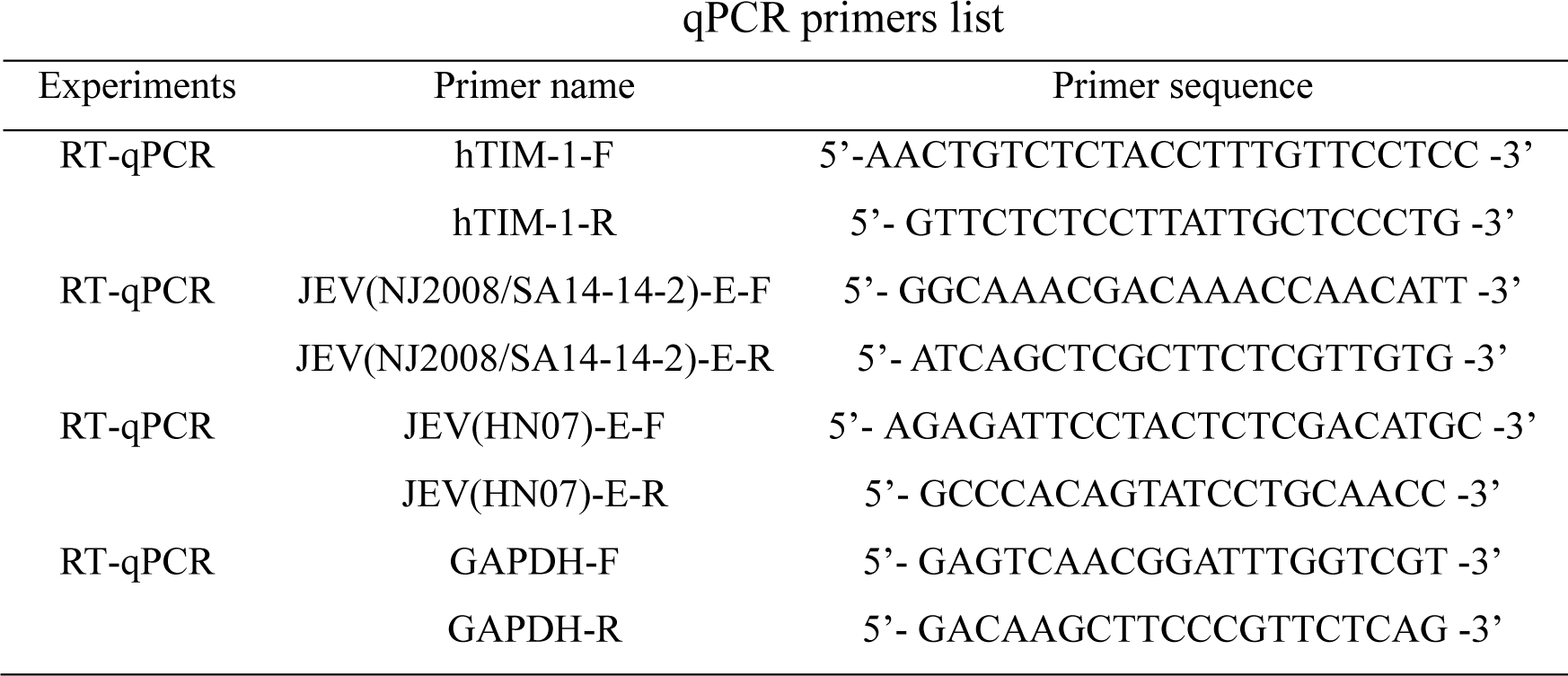

